# Differential regulation of microRNA-15a by radiation affects angiogenesis and tumor growth via modulation of acid sphingomyelinase

**DOI:** 10.1101/242933

**Authors:** Shushan Rana, Cristina Espinosa-Diez, Rebecca Ruhl, Charles R Thomas, Sudarshan Anand

## Abstract

Activation of acid sphingomyelinase (SMPD1) and the generation of ceramide is a critical regulator of apoptosis in response to cellular stress including radiation. Endothelial SMPD1 has been shown to regulate tumor responses to radiation therapy. We show here that the SMPD1 gene is regulated by a microRNA (miR), miR-15a, in endothelial cells (ECs). Standard low dose radiation (2 Gy) upregulates miR-15a and decreases SMPD1 levels. In contrast, high dose radiation (10 Gy and above) decreases miR-15a and increases SMPD1. Ectopic expression of miR-15a decreases both mRNA and protein levels of SMPD1. Mimicking the effects of high dose radiation with a miR-15a inhibitor decreases cell proliferation and increases active Caspase-3 & 7. Mechanistically, inhibition of miR-15a increases inflammatory cytokines, such as IP10, activates caspase-1 inflammasome and increases Gasdermin D, an effector of pyroptosis. Importantly, both systemic and vascular-targeted delivery of miR-15a inhibitor decreases angiogenesis and tumor growth in a CT26 murine colorectal carcinoma model. Taken together, our findings highlight a novel role for miR mediated regulation of SMPD1 during radiation responses and establish proof-of-concept that this pathway can be targeted with a miR inhibitor.

## Introduction

Technological advances such as stereotactic body radiation therapy (SBRT) and stereotactic radiosurgery (SRS) (1-5) have allowed significant improvements in therapeutic radiation dose escalation. These treatment modalities are able to ablate malignant tissue for excellent local control, however not all disease sites can be treated with these modalities due to toxicity concerns (6-10). Dose escalation does not only augments DNA damage but also involves a vast number of tumor microenvironment (TME) regulators (11). Within the TME, high dose radiation modulates the adjacent vasculature, stroma, and immune cells to contribute to the ionizing radiation (IR) response [4]. Radiation elicits endothelial cell dysfunction characterized by associated increased permeability, detachment from the underlying basement membrane, and apoptosis (12, 13). At ablative doses, greater than 8 Gy, there is rapid induction of sphingomyelinase-mediated production of ceramide, which triggers rapid onset of endothelial apoptosis (14). Indeed, it is thought that endothelial apoptosis dictates the radiosensitivity of tumors. IR-mediated cell death combined with a pro-inflammatory state contributes to an immunostimulatory profile leading to further immunogenic cell death (ICD) (15, 16).

MiRs play an important role in radiation responses of both malignant cells and the TME (17, 18). miRs are endogenous, short non-coding, single-stranded RNA spanning approximately 22 nucleotides. We and others have shown that radiation regulated miRs alter DNA damage repair pathways, pro-survival signaling pathways, cell-cycle checkpoint regulation, and apoptosis; functions which radiation therapy exploits for therapeutic gain (19-24). Our previous work identified a group of miRs regulated in the tumor vasculature in response to radiation (25). In particular, we have observed that some miRs in ECs are differentially regulated in response to different doses of radiation. We focused further attention on miRs predicted to target SMPD1. We found that miR-15a expressed the greatest magnitude difference between standard and ablative dose radiation with substantially lower miR-15a levels at higher doses. Our studies show that miR-15a targets SMPD1 in ECs and inhibition of miR-15a decreases EC and tumor cell proliferation, enhances cell death and diminishes tumor growth in a mouse CT26 colorectal carcinoma flank tumor model. Vascular-targeted nanoparticle delivery of miR-15a inhibitor is sufficient to both decrease tumor growth and angiogenesis. Consistent with the immunostimulatory role of miR-15a deficiency in autoimmune and infectious settings (26, 27), we found miR-15a inhibition increased CXCL10 expression and caspase-1 activation. In summary, our findings establish a new miR based regulatory pathway that affects SMPD1 and therefore vascular cell death in response to radiation dose. Inhibition of this pathway may mimic features of high dose radiation and therefore offers avenues for the development of targeted therapeutics.

## Materials and Methods

### miRNA profiling

RNA was extracted from HUVECs at 6h post radiation with either 2 Gy or 20 Gy and miRs were profiled using TaqMan TLDA panels for human microRNAs. miRs proposed to target miR-15a as predicted by TargetScan were further analyzed. Mean fold change after normalization to housekeeping RNA, RNU48, is depicted.

### Cell Culture and Reagents

HUVECs (Lonza) were cultured in EBM-2 media (Lonza) supplemented with 10% fetal calf serum (Hyclone). CT-26 cells (ATCC) were culture in RPMI media supplement with 10% fetal calf serum and antibiotics. HCT-116 cells (ATCC) were cultured in McCoy’s supplemented with 10% Fetal Calf Serum and antibiotics. Cells were tested and found negative for mycoplasma contamination before use in the assays described.

### Transfections

Cells were reverse transfected with miR-15a-5p mimics, inhibitors and their respective controls using Lipofectamine RNAiMAX (Invitrogen) according to manufacturer’s instructions. miR mimics and inhibitors were purchased from Life Technologies or Exiqon.

### In vivo assays

All animal work was approved by the OHSU Institutional Animal Use and Care Committee. 8–10 week old Balb/C mice purchased from Jackson Labs were injected subcutaneously with 5 × 10^5^ tumor cells in Matrigel (BD) per each flank. Tumor growth was measured with calipers, with volume computed as ½ * Length * Width^2^. Mice were randomized into groups once the average tumor volume reached 100 mm^3^, approximately 7–10 days after injection. miR inhibitors were delivered i.v. in either PBS or vascular targeted 7C1 nanoparticles every two days from randomization for a total of three doses.

### Irradiation

Cells or mice were irradiated on a Shepherd Cesium-137 irradiator at a rate of approximately1.34 cGy per minute. In tumor-targeted radiation experiments, mice were restrained in a lead shield (Brain Tree Scientific) to minimize exposure to the non-tumor areas.

### Cell Titer Glo/ Caspase Glo

Cells were transfected in a 6 well plate with miR-15a-5p mimic or inhibitor, and the corresponding controls from Exiqon (Qiagen) as previously described. Cells were transferred to a 96 well plate 16 hours post-transfection (1000 cells/well). In some studies, at 24h post-transfection the cells were irradiated with 0, 2, or 5 Gy. Cell Titer-Glo and Caspase 3/7 Glo were analyzed at 48h and 96h, according to manufacturer’s instructions.

### Statistics

All statistical analysis was performed using Excel (Microsoft) or Prism (GraphPad). Two-tailed Student’s T-test or ANOVA with post-hoc corrections was used to calculate statistical significance. P values <0.05 were considered significant.

## Results

### SMPD1 expression correlates with better overall survival in cancer

We first evaluated the expression of SMPD1 in human cancers and asked if the levels of SMPD1 correlated with overall survival (Figure 1) using the online database kmplotter. We observed that in breast and ovarian cancers, SMPD1 high patients had significantly better overall survival. In lung cancer patients, data was available for patients that only received radiation therapy. In this subset, SMPD1 high patients had almost two-fold better overall survival than patients with low SMPD1 (Figure 1C). Analysis of TCGA revealed that SMPD1 is seldom mutated or amplified suggesting transcriptional and/or post transcriptional mechanisms control the expression of SMPD1.

**Figure 1.**
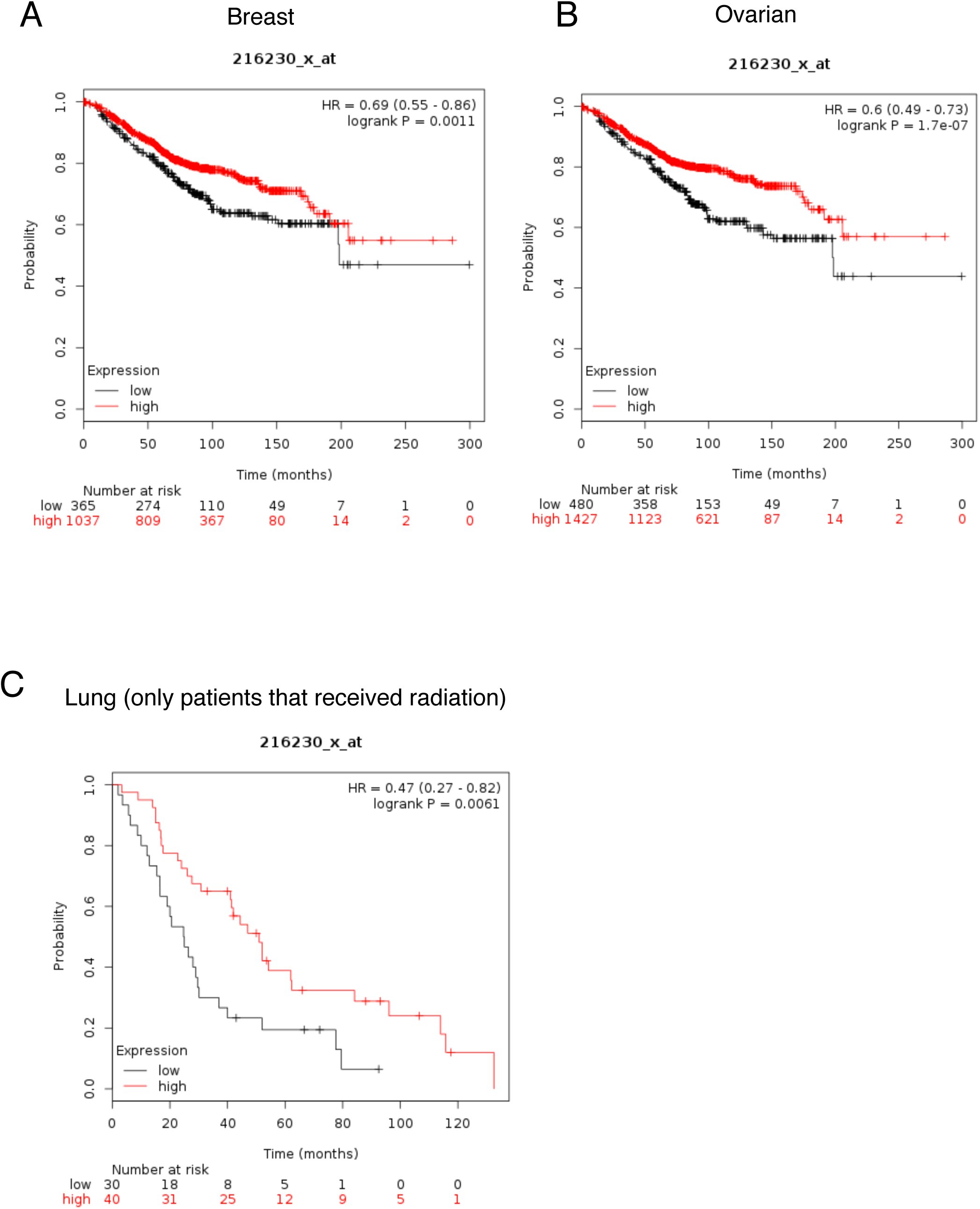
SMPD1 expression correlates with better overall survival in human cancers. Kaplan-Meier plots (kmplotter) showing overall survival in A) Breast, B) Ovarian and C) Lung cancer patients expressing high vs low SMPD1 levels. The expression levels were classified as high or low based on median expression of the gene.

### miRs regulating SMPD1 exhibit differential dose expression

Given that miRs are a major mechanism for post-transcriptional control of gene expression, we sought to identify miRs that specifically targeted SMPD1. TargetScan analysis of the SMPD1 3’ untranslated region identified miR-15 family as putative regulators of SMPD1 (Figure 2A). We chose to evaluate this using ECs as a model system since they express ~ 20 fold more SMPD1 than tumor cells. We asked if there was any miR-15a family member that was differentially regulated by radiation. HUVECs were treated with either a single 2 Gy or 20 Gy dose via Cs-137 and miRs were profiled at 6h post treatment. miR-15a exhibited the greatest differential change at 6 hours post-IR between exposure of 2 Gy and 20 Gy radiation relative to non-irradiated samples (Figure 2B). We first confirmed that endogenous miR-15a decreased at high dose radiation and the expression of SMPD1 was reciprocal to the amount of miR-15a (Figure 3A) via qRT-PCR. Subsequently, we confirmed that exogenous transfection of miR-15a significantly reduced expression of SMPD1 mRNA (Figure 3B) and protein levels (Figure 3C). These observations establish that miR-15a is differentially expressed at low vs high dose radiation and affects SMPD1 levels in ECs.

**Figure 2.**
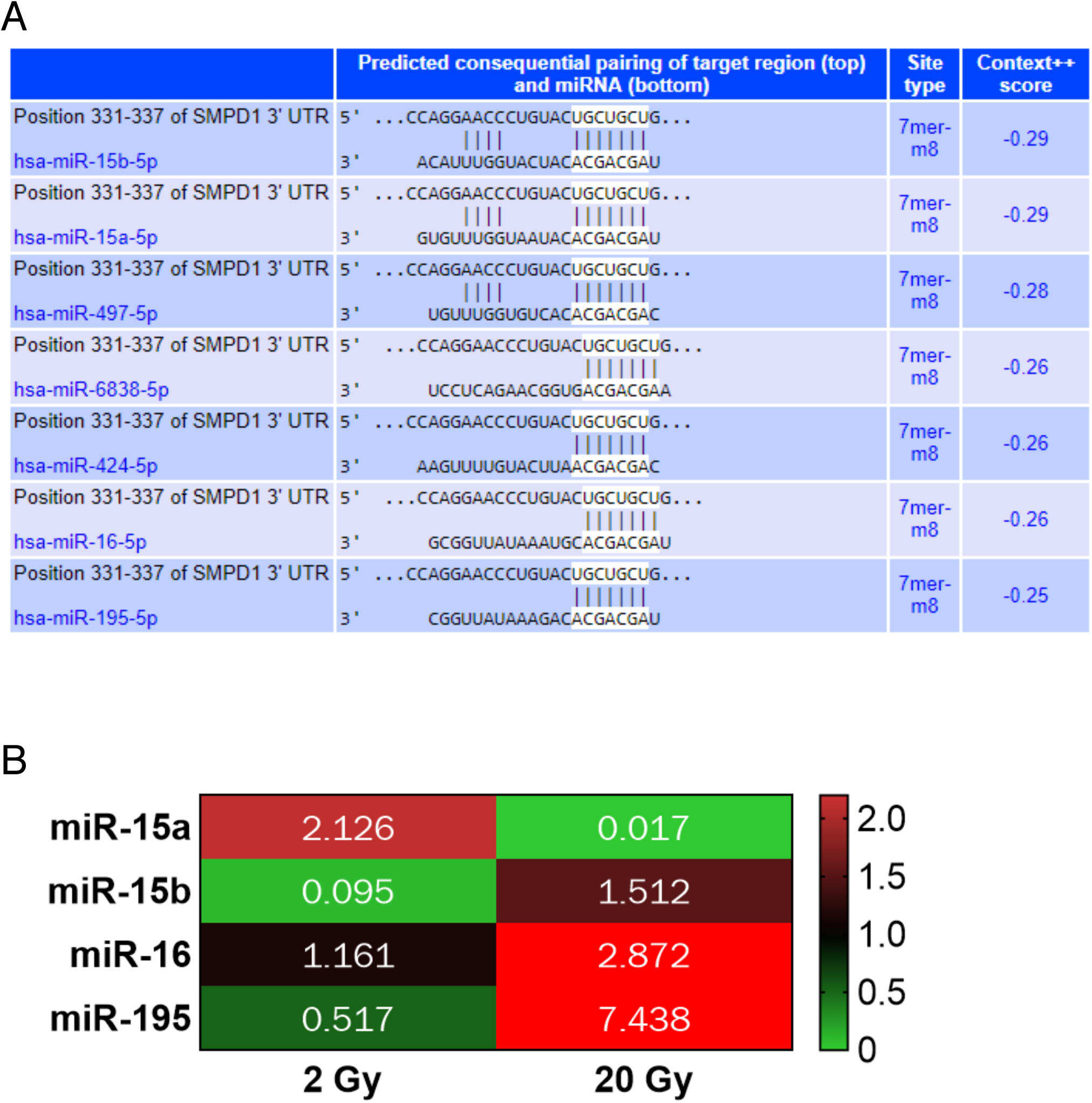
Discovery of SMPD1 targeting miRs that are differentially regulated by radiation. A) TargetScan prediction of miR candidates that harbor binding sites on the 3’ untranslated region of human SMPD1. B) miR candidates targeting SMPD1 exhibit radiation dose-dependent differential expression at 6h post-IR in HUVECs. Fold changes are indicated in colored cells relative to expression of the respective miRNA in non-irradiated samples. Red = increased expression. Green = decreased expression.

**Figure 3.**
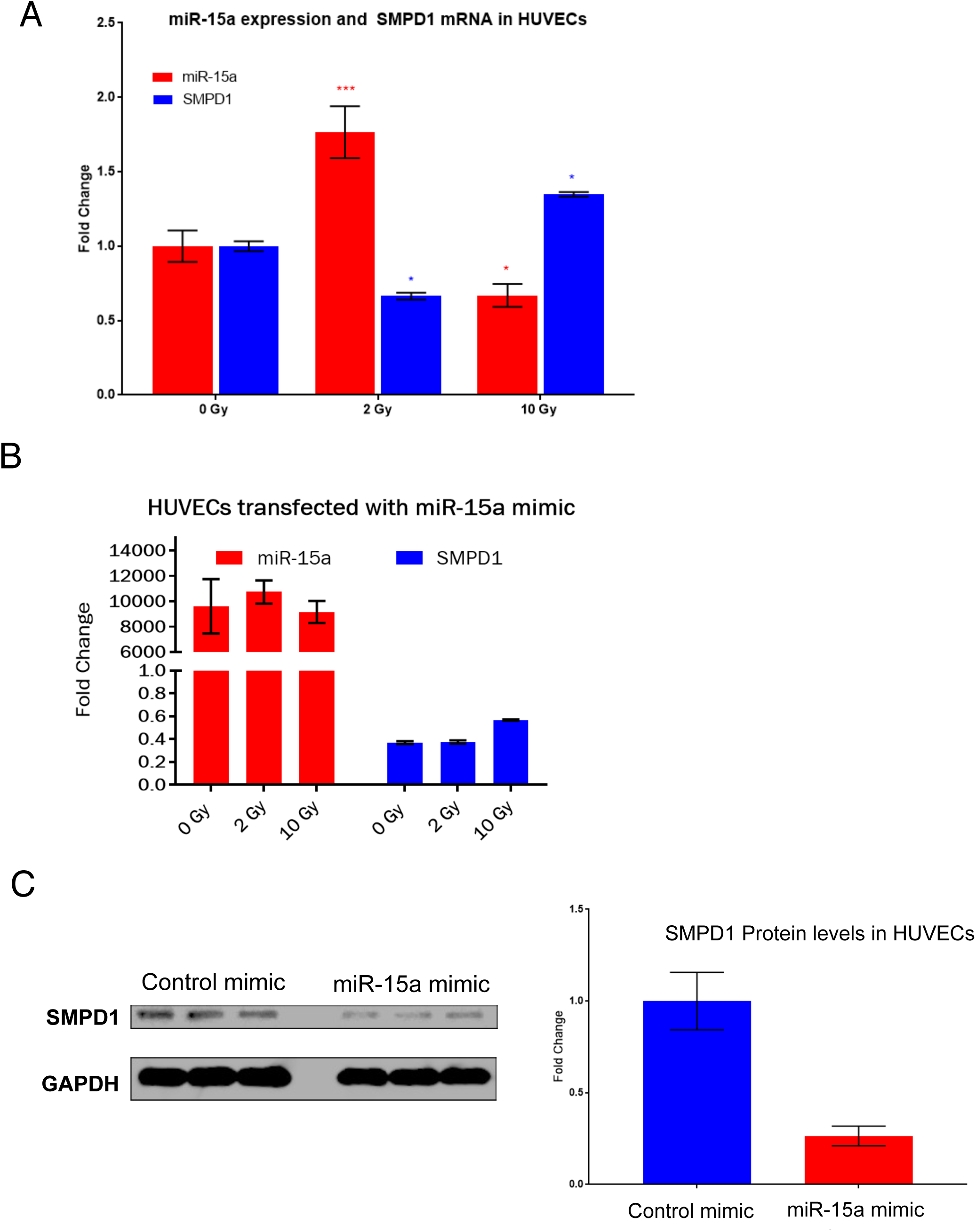
miR 15a decreases SMPD1 expression in endothelial cells. A) Reciprocal regulation of endogenous miR-15a and SMPD1 by high vs low dose radiation. HUVECs were irradiated as indicated and RNA was extracted at 18h post RT. Bars show mean ± SEM of replicates. B-C) HUVECs were transfected with either a control mimic or a miR-15a mimic. B) 24h later RNA was isolated and qRT-PCR was performed to measure the levels of miR-15a and SMPD1. C) Cells were lysed at 48h post transfection and SMPD1 protein levels were measured by western blotting. Lanes show biological replicates and bar graph shows mean band intensity ± SEM of replicates. C) *P<0.01, ***P<0.0001 per ANOVA with post hoc Dunnett’s multiple comparison test.

### miR-15a inhibition decreases HUVEC viability and increases caspase activity

Since our data indicates that high dose radiation decreased miR-15a and increased SMPD1, we asked if inhibition of miR-15a affected cell viability. HUVECs transfected with miR-15a inhibitor demonstrated dramatically decreased cell proliferation at 48h and increased Caspase activation 24h post radiation (Figure 4A-B). While noting SMPD1 is characterized by a 20 fold increased expression in ECs relative to other cell types (28), we analyzed the effects of miR-15a inhibitor on apoptosis in a similar fashion in malignant cell lines focusing on colorectal cell lines. Similar to HUVECs, miR-15a inhibitor dramatically decreased cell viability in HCT-116 cells and CT26 cells (Supplementary Figure 1). We observed that consistent with other reports, miR-15a inhibition affected inflammatory signaling in ECs by increasing IP10 (CXCL10) levels (Supplementary Figure 2A), enhancing caspase-1 inflammasome activation (Supplementary Figure 2B) and increasing the expression of Gasdermin D (Supplementary Figure 2C), a key regulator of pyroptosis (29). Pyroptosis is a lytic, regulated cell death that requires the enzymatic activity of inflammatory caspases. Since pyroptosis releases intracellular danger associated molecular patterns (DAMPs) and cytokines such as IL-β, it is thought to be a more immunogenic form of cell death (30).

**Figure 4.**
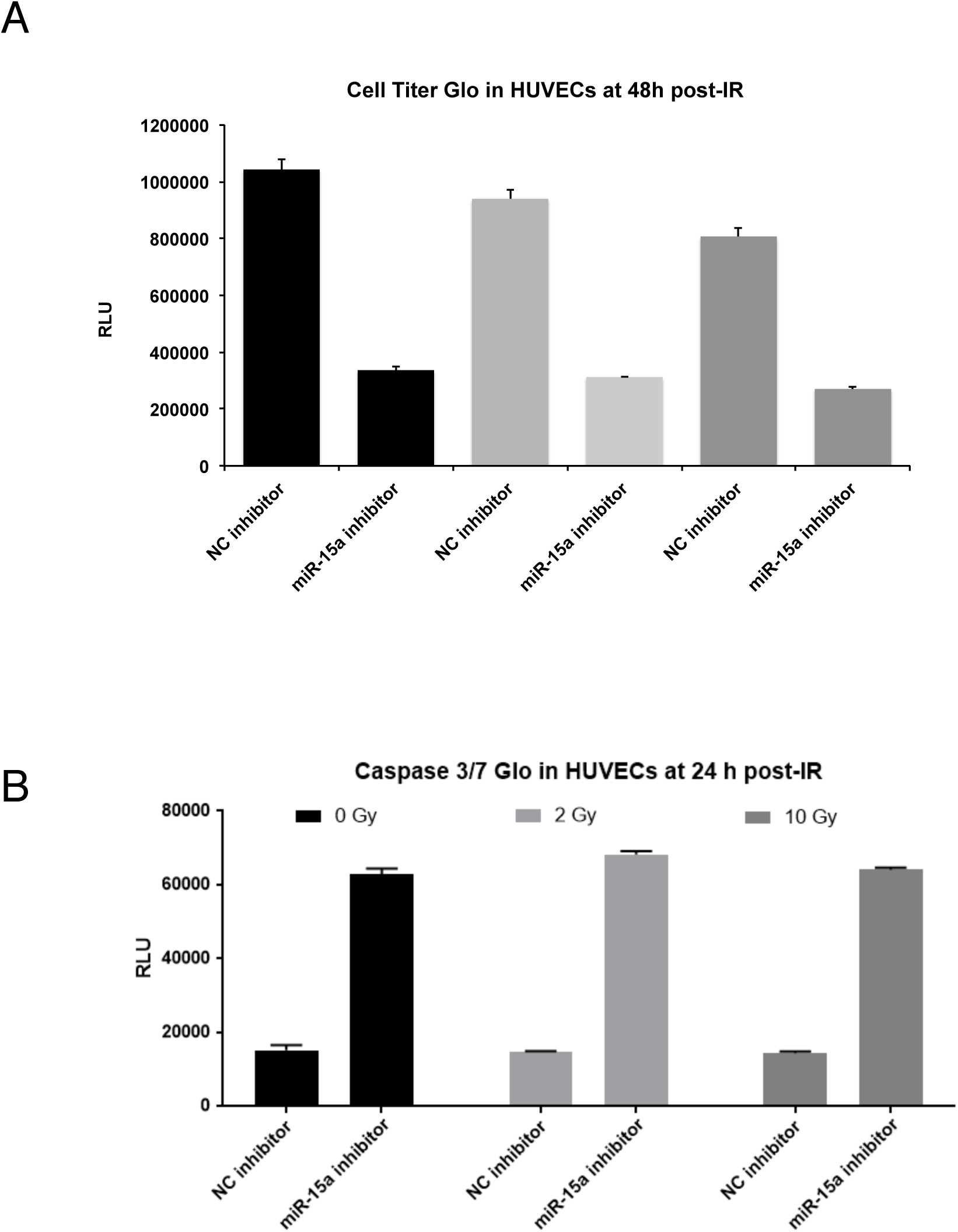
Inhibition of miR-15a decreases endothelial cell proliferation and enhances cell death. A. HUVECs were transfected with either a control negative inhibitor or a miR-15a inhibitor. 24h after transfection, cells were irradiated with either a 2 Gy or 10 Gy dose in a single fraction. 48h post radiation proliferation (A) or cell death (B) was measured using a luciferase based Cell Titer glo assay (A) or Caspase 3 & 7 CasGlo assay (B). Bars indicate means ± SEM of 3 technical replicate wells. One of at least two independent experiments is shown.

### Inhibition of miR-15a in the vasculature decreases tumor growth and angiogenesis

We next assessed whether miR-15a inhibitor had any effects on tumor growth *in vivo* and if these effects were dependent on its regulation of angiogenesis. In a murine CT26 colorectal carcinoma flank tumor model, systemic treatment with i.v. injected miR-15a inhibitor resulted in an approximately 50% decrease in tumor growth after 7 days (Figure 5A). Since our in vitro experiments demonstrated miR-15a inhibition also affected CT26 proliferation, it is possible that this tumor delay was a result of direct tumor cell inhibition. To address this, we took advantage of a vascular-targeted nanoparticle that we have established as an efficient platform for delivering miRs to tumor vasculature and not tumor cells. We found that delivery of vascular-targeted miR-15a inhibitor in the same model was sufficient to decrease tumor burden (Figure 5B). Importantly, the tumors treated with miR-15a inhibitor had a significant decrease in angiogenesis as measured by CD31 area (Figure 5C). Taken together, our observations indicate that miR-15a, a regulator of SMPD1, is inhibited by high dose radiation in ECs. A synthetic miR-15a inhibitor not only decreased EC proliferation *in vitro* but also decreased angiogenesis and tumor growth *in vivo.*

**Figure 5.**
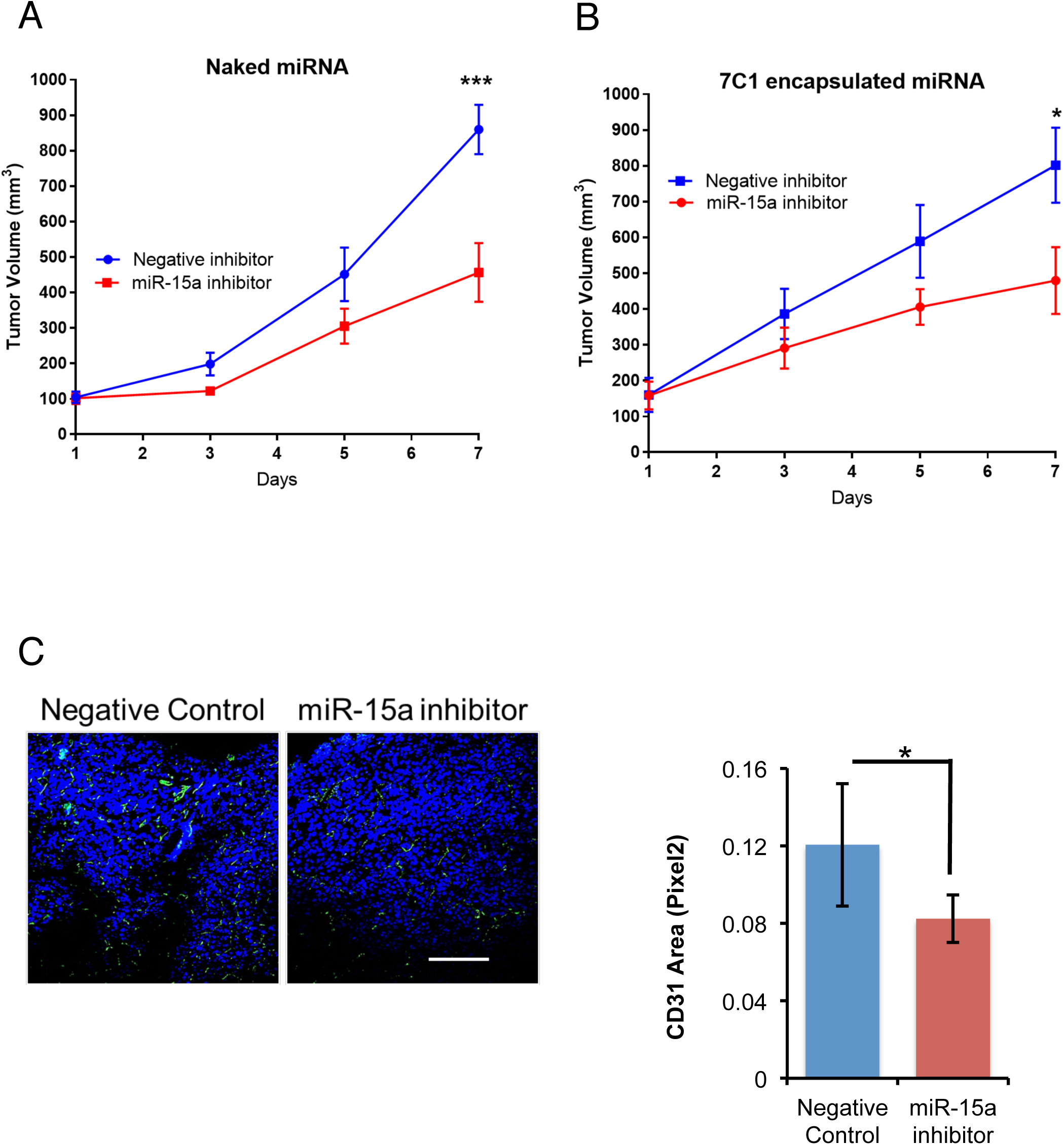
Systemic or vascular inhibition of miR-15a decreases tumor burden. A) CT26 tumors were implanted subcutaneously in Balb/C mice (N=5 mice per group, two tumors per mouse). Once tumors reached 100 mm^3^ volume, mice were randomly assigned to either a negative control inhibitor group or a miR-15a inhibitor (20mg/kg, i.v. in PBS). Mice were treated every two days for a total of three treatments. ***P<0.01; ANOVA. B) In the same model, mice were randomly assigned to receive either a negative control inhibitor or a miR-15a inhibitor in vascular-targeted 7C1 nanoparticles (1mg/kg, i.v.). * P<0.05; ANOVA. C) CD31 staining showing angiogenesis in the tumors from B). Bars show mean + SEM of 3–4 tumor sections from each mouse. *P<0.05, Student’s T-test.

## Discussion

The importance of the TME in radiation has been elucidated with the advent of new technologies and techniques allowing safer radiation dose escalation that engages the TME components (11). Kolesnick et al were among the first to demonstrate the importance of dose magnitude in eliciting rapid endothelial apoptosis via SMPD1 translocation to the plasma membrane. This translocation of SMPD1 produced ceramide thereby facilitating enhanced FAS-FASL and TNFRSF10-TNFLSF10 apoptotic signaling (28). While earlier pre-clinical models focused attention on single high dose radiation, this is not directly clinically applicable to most disease sites given dose limitations to adjacent critical organs. With this constraint, total radiation dose is divided over several days to allow sublethal damage repair of normal tissue. Using a syngeneic CT26 colorectal cancer model, Zhu et al compared fractionation between 6 Gy × 5 fractions and 12 Gy × 3 fractions. In the 6 Gy cohort, only a cumulative dose of 12 Gy or higher led to incremental increased SMPD1 activity, increased endothelial cell apoptosis, and decreased microvessel density. In contrast, multiple administrations of 12 Gy did not significantly change SMPD1 function or EC apoptosis rates (31).

As radiation dose dictates SMPD1 activity, as well as the expression of distinct miRs, we asked whether miRs with predicted binding to the SMPD1 3’-UTR also exhibited dose dependent differential expression. Interestingly, among our miRNA microarray, there were three miRs targeting SMPD1, which increased with higher doses of radiation. However, just a single miR, miR-15a was increased nearly 2-fold at 2 Gy and decreased significantly with the ablative dose radiation of 20 Gy. Recent insight into vascular miR-15a, elucidates oxidative stress as an inhibitor of miR-15a expression and the subsequent rise in SMPD1 activity. In retinal ECs, Wang et al confirmed that miR-15a binds directly to the 3’-UTR of SMPD1, and also that miR-15a inhibition significantly increases ceramide production. Indeed, miR-15a inhibition has been shown to increase expression of pro-inflammatory cytokines such as IL-6, IL-1β, and TNF-α (32) and increased leukostasis, elevated CD45, and NF-κB levels (33) in different pathophysiological contexts.

In the oncogenic context, miR-15a inhibition has been shown to enhance the innate immune response in favor of anti-tumor immunity. Yang et al (34) found that miR-15a deficiency inhibited tumor growth and prolonged survival in an orthotopic glioma model. In these experiments, they demonstrated miR-15a deficiency led to an influx of CD8+ T cells, decreased expression of inhibitory receptors including PD-1, Tim-3, and LAG-3, and increased inflammatory cytokine production.

Given the heterogeneity of cancer and versatile nature of miRs, miR-15a’s role as either an oncogenic miR or a tumor suppressive miR does not lie firmly within one category. Several cancer including non-small cell lung cancer and breast cancer, express lower miR-15a levels. This decrease in the miR has been linked to increased tumor growth and radioresistance that is reversible through miR-15a overexpression (35, 36). In colorectal cancer, a recent analysis of 182 patients found that miR-15a overexpression is associated with a worse 5-year progression free survival and overall survival (68% vs 88%, p=0.001; 60% vs 74%, p=0.035, respectively) (37). However, the dichotomic behavior is not unique to miR-15a, being a largely an oversimplified classification for this molecule able to regulate multiple targets in a context dependent fashion (38).

We chose to use colorectal cancer as our model for in vivo studies given the above findings. However, the primary focus remains the influence of vascular miR-15a on the TME to effect anti-cancer activity. SMPD1 is also a known target of miR-15a, consistent with our findings, and they both exhibit dose dependent reciprocal expression. MiR-15a inhibition decreased cellular viability, increased endothelial caspase activity and enhanced both inflammasome activation and Gasdermin expression. These mechanisms suggest that miR-15a inhibition maybe potent due to its ability to drive pyroptosis in the vasculature, which would be beneficial in a therapeutic context. On the basis of these observations, we propose that inhibition of miR-15a offers a unique approach to suppress tumor growth.

## Acknowledgements

We thank Dr. Liana Tsikitis (OHSU) for useful discussions. We thank Drs Daniel G Anderson and Omar F. Khan (MIT) for 7C1 nanoparticles. We thank LaTroy Robinson for technical help. We acknowledge the OHSU Advanced Light Microscopy Core, Knight Cancer Institute Flow Cytometry Core and the Gene Profiling Shared Resource for technical help and useful discussions. This work was supported by US NIH grant R00HL112962, R56HL137779 and an innovative research grant from the American Heart Association (17IRG33400218) to S.A and a seed grant from ASTRO to S.R (Grant ID 534775).

